# AlphaFlex: Accuracy modeling of protein multiple conformations via predicted flexible residues

**DOI:** 10.1101/2025.07.11.664327

**Authors:** Lingyu Ge, Xinyue Cui, Kailong Zhao, Xiaogen Zhou, Yang Zhang, Guijun Zhang

## Abstract

Understanding protein conformational dynamics is critical for elucidating disease mechanisms and facilitating structure-based drug design. While AlphaFold2-based methods have advanced static structure prediction, modeling multiple conformation states with both accuracy and efficiency remains challenging. We present AlphaFlex, an innovative framework that integrates flexible residue prediction with a directed masking strategy for multiple sequence alignments. By selectively relaxing co-evolutionary constraints in dynamic regions, AlphaFlex enables accurate prediction of biologically relevant states while preserving structural fidelity. Benchmarking on 69 apo-holo pairs from the CoDNaS dataset, including enzymes, binding proteins, and chaperones, reveals AlphaFlex’s superior performance in both flexible residue identification and multiple conformation prediction. Comparative analysis against AF-Cluster, AF2-conformations, and AlphaFlow demonstrates AlphaFlex’s significant advantage, achieving the highest success rate (42%) in accurately predicting apo/holo structures (TM-score >0.95 for both states) while consistently outperforming these methods in reproducing both apo and holo conformations. Notably, AlphaFlex maintains robust performance for membrane proteins, resolving inward- and outward-facing conformations with equal fidelity. Case studies further reveal its unique ability to sample physically plausible intermediate states along transition pathways. These results establish AlphaFlex as a pivotal computational tool for mapping protein conformational landscapes, with promising potential to accelerate structure-based therapeutic development.

## Introduction

Proteins serve as the fundamental molecular machinery of life, forming the structural basis of cellular organization and orchestrating virtually all biological processes. Deciphering their three-dimensional structures is essential for understanding functional mechanisms in both physiological and pathological contexts^1^. Recent transformative advances in deep learning-driven protein structure prediction, exemplified by tools such as AlphaFold^2,3^, RoseTTAFold^4^, and ESMFold^5^, have effectively resolved the long-standing challenge of accurately predicting static monomeric protein structures. Notably, the biological functionality of proteins is intrinsically governed by their dynamic nature and remarkable capacity to adopt multiple conformational states during crucial processes, including enzymatic catalysis, allosteric regulation, and cellular signal transduction^6^. While experimental techniques, including X-ray crystallography^7^, nuclear magnetic resonance^8^ (NMR), and cryo-electron microscopy^9^ (cryo-EM), can provide high-resolution structural information, they face inherent limitations in characterizing dynamic conformations, such as insufficient temporal resolution, structural averaging effects, and substantial experimental demands^10,11^. The remarkable progress achieved in static protein structure prediction stands in sharp contrast to the persistent experimental challenges in resolving dynamic conformational ensembles, propelling computational prediction of protein dynamics to the forefront of structural biology research. Addressing this critical challenge holds the promise of bridging the gap between dynamic conformation determination and functional interpretation, while simultaneously providing unprecedented insights into the mechanistic foundations of biological processes at the molecular level.

Computational approaches for predicting protein multiple conformations have evolved into three principal methodologies, each with distinct advantages and inherent limitations. The field is currently dominated by molecular dynamics (MD) simulations, Monte Carlo (MC) sampling techniques, and artificial intelligence (AI)-based generative models. MD simulations employing physics-based force fields deliver atomic-resolution trajectories, with deep learning accelerators such as AI^2^BMD^12^ having substantially improved their computational efficiency. Nevertheless, these simulations remain constrained by the immense dimensionality of conformational space and kinetic barriers that limit sampling of biologically relevant timescales^13^. MC sampling methods, which explore conformational space through energy-guided Markov chains, continue to grapple with dual challenges: imperfect energy functions that fail to capture subtle interactions and inherent sampling inefficiencies. While innovations like MultiSFold^14^ and M-SADA^15^ have enhanced conformational coverage through advanced sampling strategies, persistent difficulties in overcoming energy barriers and minimizing sampling bias underscore the need for improved algorithms and refined energy models. AI-based approaches have established generative modeling as a leading paradigm for protein multiple conformations prediction. Several innovative approaches exemplify this trend. AlphaFLOW^16^ has developed a diffusion-based framework incorporating flow matching techniques to generate diverse conformations through refined AlphaFold architectures. In parallel, idpGAN^17^ has implemented generative adversarial networks for coarse-grained conformational set prediction, while DiG^18^ has successfully combined deep learning with simulated annealing algorithms to model equilibrium distributions. Generally, these promising methodologies share a fundamental limitation is the scarcity of experimentally validated multiple conformational data, which fundamentally restricts their training and validation. This data deficiency represents the most significant barrier to developing robust, generalizable models for protein dynamics prediction.

The revolutionary breakthrough of AlphaFold2^2^ in static protein structure prediction has laid a vital foundation for the development of multiple conformation prediction methods. The core scientific rationale for this extension stems from its unique capability to decode dynamic information: Firstly, the co-evolutionary information within multiple sequence alignments (MSA) not only encodes static structural details but also reveals distinct functional states by capturing the correlated evolutionary patterns of allosterically coupled residues. Secondly, its iterative refinement mechanism for structural modules inherently achieves directed relaxation of conformations across the energy landscape. Empirical studies from the recent CASP15 and CASP16 competition demonstrate that methods based on the enhanced AlphaFold2 framework, through optimized MSA processing strategies and the incorporation of enhanced sampling techniques, now represent the most effective technical approach for predicting multiple functional protein conformations^19^.

Enhanced sampling methods strategically perturb MSA to modulate co-evolutionary information in AlphaFold2 inputs, with representative approaches including AF-Cluster^20^ leveraging MSA similarity clustering to capture distinct conformational states, SPEACH_AF^21^ employing random alanine substitution to weaken co-evolutionary constraints, AF2_conformations^22^ and Subsampled AF2^23^ utilizing random subsampling with reduced MSA depth to generate conformational ensembles, and AFsample2^24^ applying randomized column masking at specific ratios to attenuate co-evolutionary informations. Beyond direct manipulation of MSA, complementary strategies leverage external structural guidance, such as the experimental distance constraints utilized in AlphaLink^25^, or implement redesigned entropy-based loss functions as demonstrated in EGF^26^ to address AlphaFold2’s inherent single conformation prediction bias. While these efforts have demonstrated success in various cases, significant challenges remain, including generally low success rates, the need to generate large numbers of conformational decoys to identify a few biologically relevant states, and target-dependent accuracy variations reflecting systematic biases in current energy landscape sampling. Most critically, the brute-force enumeration paradigm of existing perturbation strategies results in severely compromised algorithmic efficiency. This acute imbalance between computational cost and benefit urgently demands the development of new prediction methods that combine energy landscape-aware directed sampling algorithms with well-defined, effective MSA manipulation protocols to enable precise capture of functional conformations, rather than relying on computationally intensive random exploration.

Here, we present AlphaFlex, an MSA optimization strategy guided by flexible residues exhibiting high dynamics (residue displacements ≥ 2 Å), for predicting protein multiple conformational states. This approach builds on the key insight that co-evolutionary information within MSAs encodes not only static structural constraints but also dynamic conformational information, where specific residue pairs demonstrate significant correlations with conformational transitions. AlphaFlex employs a deep learning model, trained on a large dataset of experimental structures and MD simulation trajectories (see the **Methods** section for data details), to accurately predict residue-level flexibility distributions, which are then further used for targeted masking of corresponding MSA columns to selectively attenuate dominant conformation signals while enhancing minor conformation features, preserving evolutionary constraints in structural core regions. This approach establishes a quantitative relationship among structural dynamics, evolutionary features, and functional conformations, achieving robust multiple conformations prediction while fully retaining AlphaFold2’s original architectural framework. Using an independent test set of 69 apo/holo state proteins from CoDNaS^27^, we demonstrate that AlphaFlex achieves significant improvements in both flexible residue prediction accuracy and efficient generation of multiple conformations compared to existing methods.

## Results

### AlphaFlex overview

AlphaFlex is a two-stage computational framework for predicting multiple conformational states from a single protein structure (**Fig. 1**). In the initial protein flexible residue prediction (**Fig. 1a**), the system extracts three complementary feature categories: (1) structural features comprising geometric descriptors (ultrafast shape recognition^28^ (USR), voxelization^29^, secondary structures, distance and orientation maps), evolutionary conservation metrics (BLOSUM62), and physicochemical properties (Rosetta^30^ one-body and two-body statistical potentials); (2) MSA Transformer embeddings that capture co-evolutionary patterns; and (3) structural profile features derived from structural alignments against the AlphaFold Database (AFDB)^31^. Notably, the structural profile features quantitatively characterize conformational variations between target and remote homologous structures, directly capturing evolutionarily conserved dynamic properties since homologous proteins typically exhibit conserved conformational dynamics. These multimodal features are then integrated and processed through an attention-based and deep residual network-based^32^ encoder-decoder, which predicts residue-specific flexibility probabilities on a continuous scale from 0 (rigid) to 1 (fully flexible). Complete feature descriptions are provided in **Supplementary Table 1**. In the next multiple conformation sampling stage (**Fig. 1b**), the predicted residue-specific flexibility probabilities direct a novel MSA masking strategy (Specific strategies are detailed in the “**Methods**” section.) that selectively weakens co-evolutionary constraints in dynamic regions. Combined with deep subsampling, this generates diverse sub-MSAs capturing distinct conformational states, which are then processed through AlphaFold2^1^ to produce the final structural models.

**Fig. 1.**
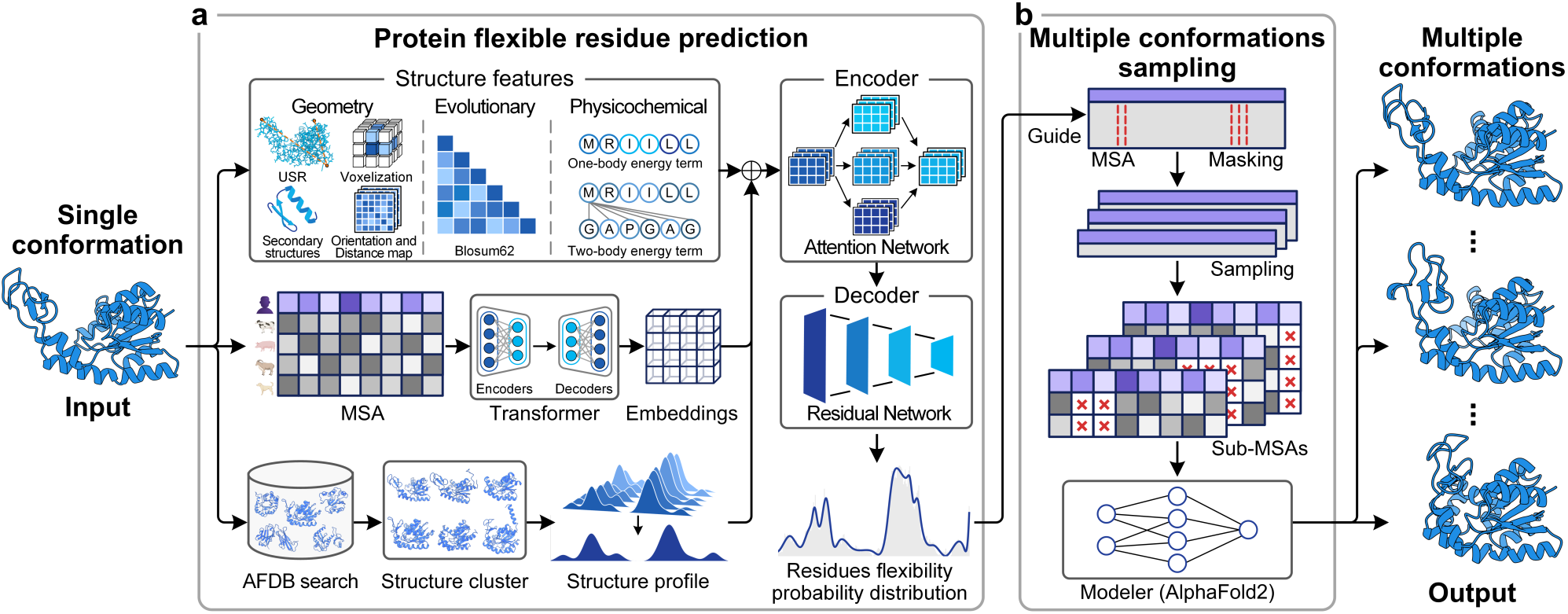
Overview of AlphaFlex. AlphaFlex predicted multiple protein conformations, using protein structure from the Protein Data Bank (PDB)^33^ or predicted models like AlphaFold2 as input, and aims to predict multiple conformations. **a,** Protein flexible residue prediction. This stage predicts per-residue flexibility using a deep learning model. The model extracts three complementary feature categories, including structural features (geometry, evolutionary, and physicochemical), MSA Transformer embeddings, and structure profile features, processed through attention and deep residual networks to produce a residue flexibility probability distribution. **b,** Multiple conformations sampling. In the second stage, the predicted flexibility distribution guides the MSA masking process. The masked MSA then undergoes a deep sampling procedure to generate multiple distinct sub-MSAs. Each of these sub-MSAs is subsequently fed into AlphaFold2 to predict diverse protein conformations.

AlphaFlex addresses a fundamental limitation in protein structure prediction where conventional methods (e.g., AlphaFold-Multimer^34^) systematically bias toward static, lowest-energy conformations due to their inability to disentangle conformation-dependent co-evolutionary information^35^. To overcome this challenge, AlphaFlex introduces an innovative “flexibility-guided MSA processing” strategy that first applies flexible residue-specific masking to MSAs, selectively attenuating non-target conformational noise while preserving state-specific evolutionary information. This is followed by deep subsampling to generate conformationally distinct sub-MSAs, which are processed through AlphaFold2 to produce structurally diverse, biophysically plausible multiple conformations. Benchmarking on a test set of 69 apo/holo proteins demonstrates that AlphaFlex not only improves success rates for multiple conformations prediction by 16.0% compared to the state-of-the-art method (we define success as modeling TM-score^36^ accuracy greater than 0.95 for both apo and holo states), but also enhances single-state accuracy (reducing RMSD by 5.1% for apo states and 5.3% for holo states), providing a robust solution for predicting functional conformational transitions from static structures.

### Performance on apo/holo multiple conformations prediction

To comprehensively evaluate the multiple conformations modeling capacity of AlphaFlex, we initially selected 91 apo/holo protein pairs from CoDNaS^27^ as the candidate test set. The training set was derived from PDB (January 2024 release) through sequential redundancy reduction: primary clustering at 80% sequence identity generated 65,327 clusters, followed by centroid screening at 30% identity to yield 20,932 non-redundant clusters. By applying strict cross-validation at a 30% sequence identity threshold between training and test sets, we established 17,485 independent training clusters and a final evaluation set of 69 stringently curated apo/holo pairs, ensuring an unbiased assessment of modeling performance (see **Methods** for data construction criteria).

The benchmark test set encompassed functionally diverse protein classes, including enzymes, binding proteins, and chaperones, representing a broad spectrum of biologically relevant conformational transitions. These range from localized loop rearrangements to substantial domain movements upon ligand binding, thereby comprehensively evaluating AlphaFlex’s ability to capture both subtle and dramatic conformational changes. We systematically evaluated the performance of AlphaFlex against the baseline methods AF-Cluster^20^, AF2_conformations^22^, and AlphaFLOW^16^. To ensure a rigorous and fair comparison, each baseline method was used to generate 1,000 conformations per protein target (see **Methods** for details). For performance evaluation, we employed two distinct metrics: first, single conformation accuracy was evaluated through Root Mean Square Deviation (RMSD) and Template Modeling score (TM-score)^36^ measurements for both apo and holo states; second, the success rate was established using a stringent dual-conformation threshold requiring TM-score > 0.95 for both states, thereby assessing the model’s capacity to accurately predict conformational transitions. Detailed descriptions of these metrics are provided in **Supplementary Note 1.**

We found that AlphaFlex outperforms existing methods in both multiple conformation prediction success rates and single conformation accuracy (**Fig. 2**). Specifically, AlphaFlex achieved a 42.0% success rate (**Fig. 2a**), with both predicted apo and holo structures exceeding a TM-score greater than 0.95 across our benchmark targets in these successful cases. Among these success cases, 58.6% of the protein pairs exhibited a TM-score less than 0.95 between the apo and holo states. This represents a 16% substantial improvement over the best baseline method. Notably, AlphaFlex achieved an average RMSD of 2.008 Å (apo) and 1.471 Å (holo), demonstrating significant enhancement in accuracy compared to baseline methods, including AF-Cluster (2.975 Å/2.487 Å), AF2_conformations (2.117 Å/1.554 Å), and AlphaFLOW (2.079 Å/2.126 Å) for apo/holo states (**Fig. 2b**). This superior performance is further reflected in TM-score metrics for overall structural prediction similarity. The detailed evaluation results are listed in **Supplementary Table 2**. To further evaluate AlphaFlex’s single conformational precision accuracy, we analyzed the distribution of its predicted apo and holo conformations across various RMSD intervals (**Fig. 2c**). AlphaFlex produced 2.4% more models satisfying the high-accuracy threshold (RMSD < 2.0 Å) in both apo/holo states than the best baseline method, demonstrating its ability to generate precise multiple conformation states.

**Fig. 2.**
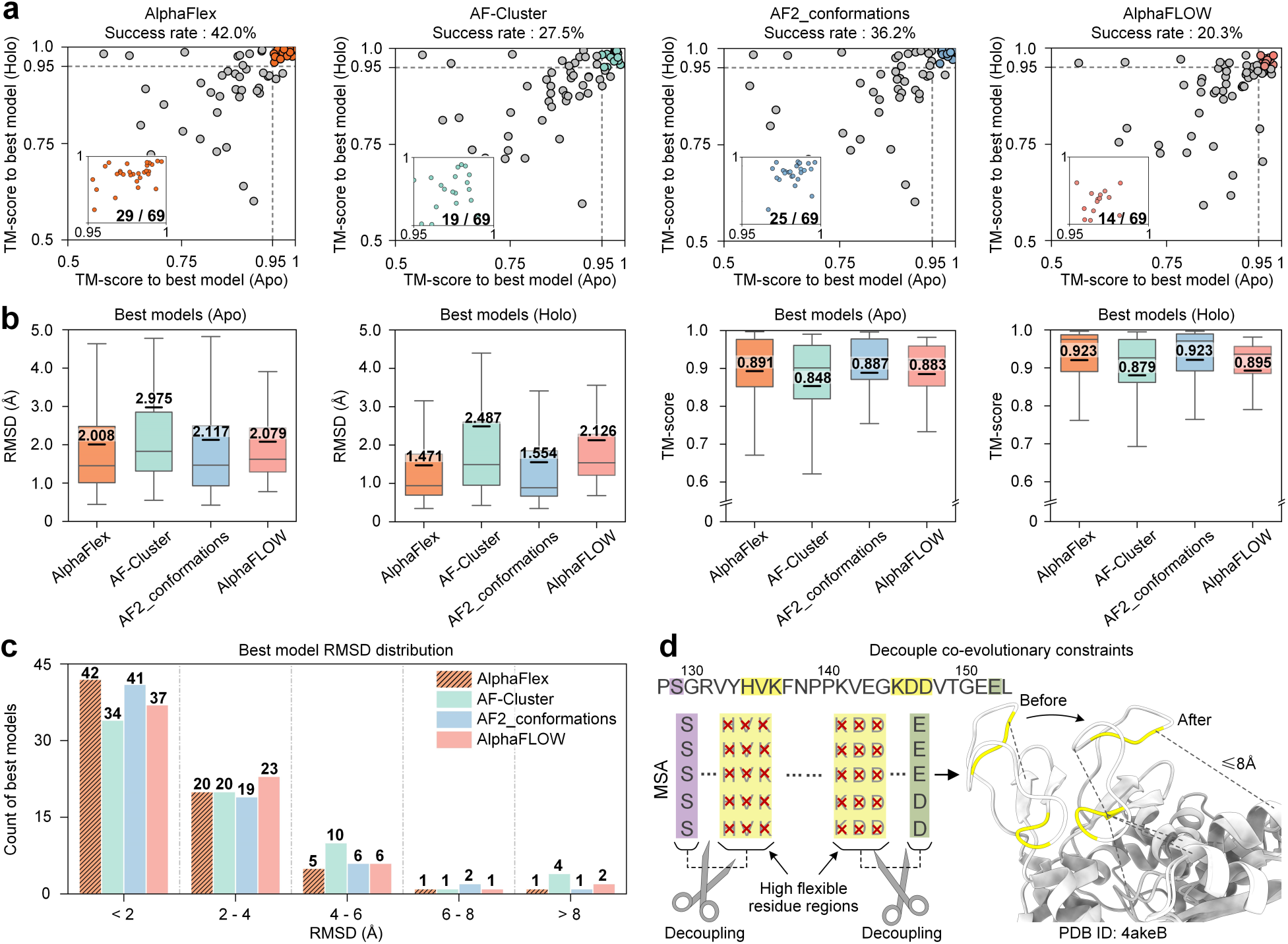
Results of protein apo/holo multiple conformations prediction. **a**, Success rates of AlphaFlex, AF-Cluster, AF2_conformations, and AlphaFLOW in predicting both apo and holo states with TM-score > 0.95. **b,** Average RMSD and TM-score for 69 protein targets with multiple conformations, predicted by AlphaFlex, AF-Cluster, AF2_conformations, and AlphaFLOW. **c,** Distribution of RMSD for the best predicted apo and holo models by different methods. **d,** Schematic diagram of AlphaFlex’s mechanism for decoupling co-evolutionary constraints between high residues probability flexible regions and conserved regions.

The consistent performance across both success rates and single conformation accuracy highlights the effectiveness of AlphaFlex’s flexibility-guided MSA processing strategy. This strategy selectively masks MSA columns corresponding to residues with high probability flexibility (**Fig. 2d**), thereby attenuating the strong co-evolutionary information that typically links these dynamic regions to more conserved segments. In unprocessed MSA, such coupling may bias the model’s prediction toward a single, low-energy conformation. By contrast, AlphaFlex decouples these co-evolutionary constraints within MSA, effectively releasing flexible regions from the constraints of conserved co-evolutionary patterns. This relaxation mechanism enables the model to sample from a broader conformational space, allowing it to generate alternative states that were suppressed during inference. The above experimental results confirm that AlphaFlex significantly enhances protein structure prediction by effectively discriminating between conformation-specific coevolutionary signatures and background evolutionary constraints. This critical capability enables accurate prediction of multiple biologically relevant conformational states while maintaining high structural fidelity.

### Performance on membrane protein multiple conformations prediction

Membrane proteins play important roles in biological processes such as signal transduction and substrate transport. However, the inherent complexity of their conformational changes poses a major challenge to the modeling of multiple conformations^37^. To further examine whether AlphaFlex can capture both inward-facing and outward-facing conformations, we assessed its performance on a non-redundant test set^38^ consisting of five representative membrane proteins. AlphaFlex achieved average RMSD values to 2.716 Å (inward-facing) and 3.083 Å (outward-facing) (**Fig. 3a**), representing 7.7% and 9.0% accuracy improved over the best baseline method, respectively, and outperformed all comparison methods in TM-score evaluations (**Fig. 3b**). The detailed evaluation results are listed in **Supplementary Table 3**. By analyzing the distribution of the prediction results of different methods under different RMSD thresholds (**Fig. 3c**), we found that the proposed method can predict 3/5 inward-facing conformations and 2/5 outward-facing conformations with high precision (RMSD < 2.0 Å). These results demonstrate that AlphaFlex possesses the capability to capture the two key conformational states of membrane proteins during biological function.

**Fig. 3.**
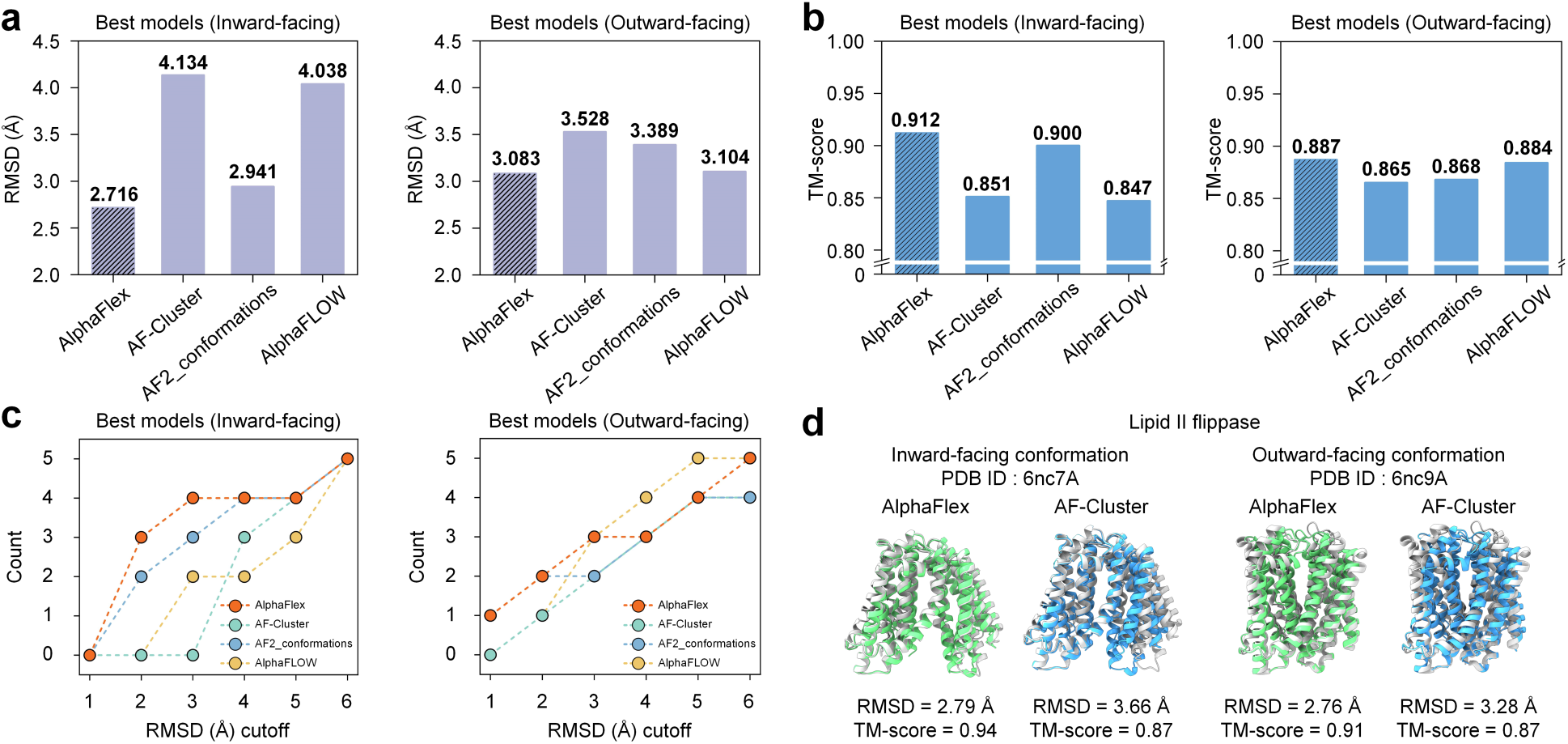
Results of membrane protein multiple conformations prediction. **a**, Average RMSD of the best models for inward-facing and outward-facing conformation, comparing different prediction methods. **b,** Average TM-score of the best models for inward-facing and outward-facing conformation, comparing different prediction methods. **c,** Number of predicted conformations under different RMSD cutoffs for each method. **d,** The structural comparison of the *Lipid II flippase* (6nc7A, inward-facing; 6nc9A, outward-facing) shows experimental structures and predictions by AlphaFlex and AF-Cluster.

A notable example is the Lipid II flippase (6nc7A, inward-facing; 6nc9A, outward-facing). As illustrated in **Fig. 3d**, although AF-Cluster achieved TM-score above 0.8 for both conformational states, there is still a certain deviation in the domain orientation. To further improve these results, AlphaFlex leverages predicted residue flexibility probability distributions to dynamically adjust the co-evolutionary information embedded in the MSA, thereby enhancing its ability to explore different conformational states. As a result, it accurately predicts both inward- and outward-facing functional conformations, achieving TM-score improvements of 8.0% and 4.6% over AF-Cluster, respectively. These results highlight that directionally decoupling the co-evolutionary information within the MSA can effectively guide the model to explore a broader conformational landscape, thereby enabling more accurate multiple conformation predictions.

### Performance on flexible residue prediction

Understanding protein residue flexibility is crucial for molecular biology and biotechnological applications. From a functional perspective, highly flexible regions are often central to a protein’s conformational changes, which in turn affect their biological functions. Therefore, we aim to capture these potential conformational change regions from the structure. In this section, we defined residues as flexible if their Cα atomic displacement exceeded 2 Å during conformational changes. To evaluate AlphaFlex’s ability to predict flexible residues, we used the same 69 apo and holo protein pairs as the benchmark set for the multiple conformation prediction task. For baseline methods, including AF-Cluster^20^, AF2_conformations^22^, and AlphaFLOW^16^, flexible residues were determined based on whether the Cα distance between the two structure with the largest conformational difference exceeds 2 Å. The performance was evaluated using four metrics: the area under the receiver operating characteristic curve (ROC-AUC), accuracy, precision, and F1 score. The comparative results are summarized in **Fig. 4a**. Detailed evaluation results are provided in **Supplementary Table 4**. On average, AlphaFlex achieved a ROC-AUC of 0.810, an accuracy of 0.742, a precision of 0.491, and an F1 score of 0.521, representing improvements of 23.1%, 3.1%, 11.1%, and 22.9%, respectively, over the best baseline method. The superior performance of AlphaFlex in predicting flexible residues can be primarily attributed to its diverse training dataset, which includes multiple conformational structures obtained through experimental techniques such as X-ray crystallography, NMR, and cryo-EM. Furthermore, subsequent fine-tuning on MD simulation data further improved the model’s capacity to capture realistic dynamic behavior.

**Fig. 4.**
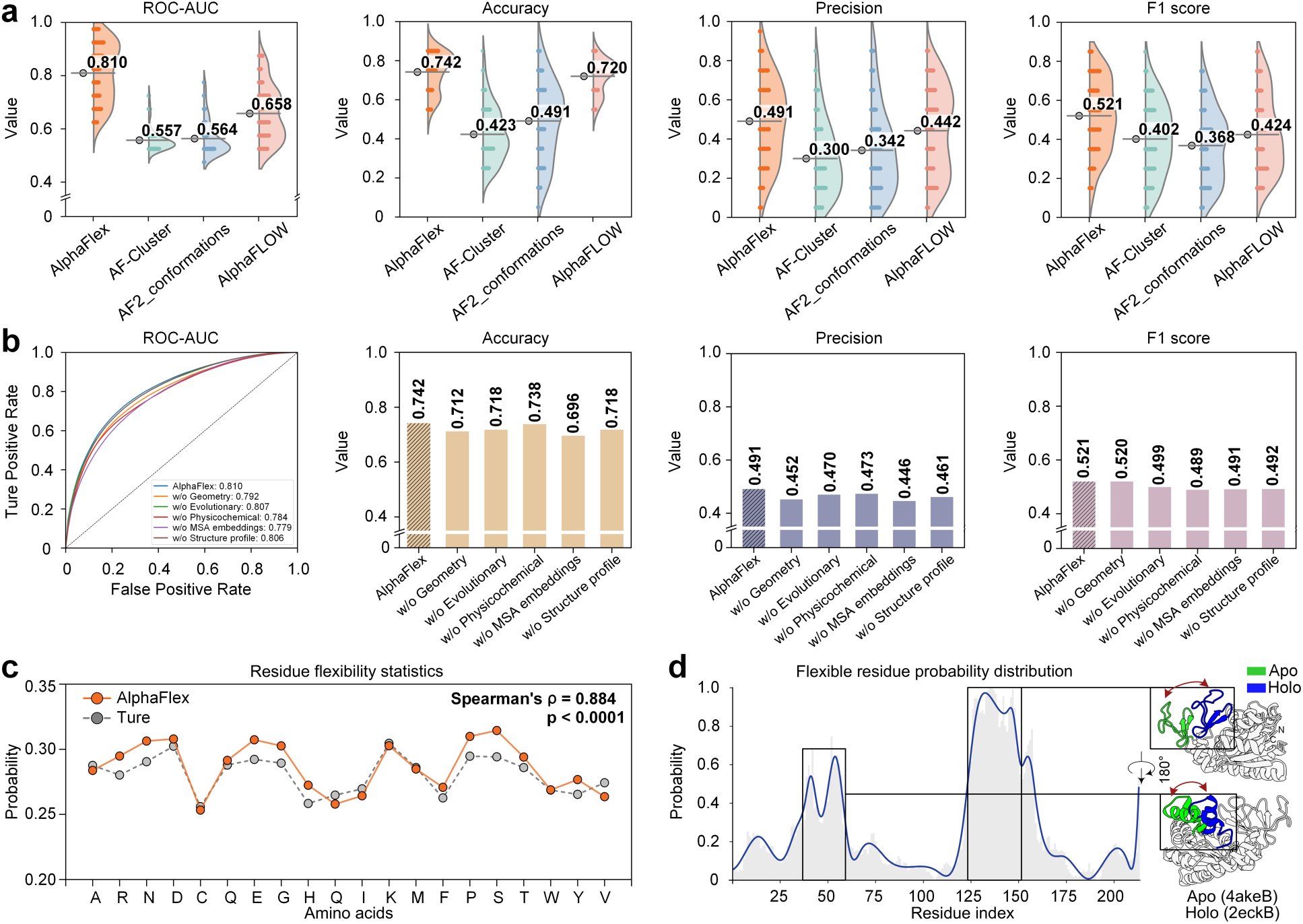
Results of the protein flexible residue prediction. **a**, Experimental results of protein flexible residue prediction. **b,** Ablation study demonstrating the impact of removing different feature types on the performance of AlphaFlex in flexible residue. **c,** Statistical analysis of the flexibility propensity of different amino acid types. **d,** Schematic diagram illustrating AlphaFlex’s predicted probability distribution of flexible residues and corresponding structural changes.

To verify the contribution of different feature sets to the model’s performance, we conducted an ablation study by systematically removing one feature type at a time. This resulted in five model variants, named as w/o Geometry, w/o Evolutionary, w/o Physicochemical, w/o MSA embeddings, and w/o Structure profile. We evaluated these variants against the full model using ROC-AUC, Accuracy, Precision, and F1 score. As summarized in **Fig. 4b** and detailed in **Supplementary Table 5**, these results collectively demonstrate that all feature components are crucial for the model’s predictive performance. The structural features (geometry, Evolutionary, and physicochemical) contribute rich structural and biochemical information, which is essential for capturing residue-level conformational dynamics. Complete feature descriptions are provided in **Supplementary Table 1**.

Our model integrates a rich array of multi-source features to comprehensively capture protein behavior. Geometric features incorporate multiple aspects of spatial information, including topological relationships from Ultra-fast Shape Recognition^28^ (USR), Voxelization^29^ representations based on backbone atoms, secondary structure annotations, residue-level distance, and contact maps. These features collectively encode the conformational space and spatial organization of proteins. In parallel, evolutionary features such as BLOSUM62 capture residue conservation patterns reflecting the amino acid preferences shaped by evolutionary processes across homologous sequences. The conservation of a residue during evolution usually reflects its functional importance and may be related to its flexibility characteristics. Physicochemical features enrich the model with biochemical context by encoding residue interaction tendencies through energy-based descriptors, including hydrogen bonding, van der Waals interactions, electrostatics, and solvent accessibility. These interactions are closely associated with local conformational dynamics and flexibility. Together, these residue-level descriptors offer a comprehensive and detailed representation of protein behavior that is essential for accurate flexibility prediction.

Our ablation studies demonstrate that each feature category contributes unique and indispensable information, collectively enhancing the model’s predictive accuracy for residue flexibility. Notably, MSA embeddings showed the strongest impact on model performance. Their removal led to the most substantial performance degradation across all evaluation metrics, reducing the F1 score by 5.8% (**Fig. 4b**). This highlights the critical role of the deep co-evolutionary coupling information derived from homologous sequences, which captures long-range dependencies between residues essential for predicting flexibility. Furthermore, the structural profile contributes positively to the model’s performance, and when these features are removed, the F1 score of the model decreases by 5.6%. This is likely because structural profile features effectively capture the protein’s spatial fluctuations by integrating diverse structural homology information. Our ablation study confirms that each feature category provides unique and indispensable information, directly contributing to the model’s ability to predict residue flexibility. The integration of these diverse features allows the model to form a comprehensive biological representation, which is key to its enhanced predictive accuracy for residue flexibility.

To further investigate whether AlphaFlex captures intrinsic flexibility patterns encoded in amino acids. We compared AlphaFlex’s predicted flexibility propensities for the 20 common amino acid types, based on 69 apo/holo protein pairs (**Fig. 4c**). Spearman rank correlation analysis revealed a strong positive correlation (ρ = 0.884), demonstrating AlphaFlex’s accurate characterization of intrinsic amino acid flexibility variations. Among all types, we found that cysteine tends to exhibit the lowest predicted flexibility. This may be due to its unique ability to form disulfide bonds, which rigidify the local structure and contribute to the stabilization of protein tertiary conformations, thereby limiting large-scale conformational transitions. To provide a representative case of flexibility residues, we analyzed the *E. coli adenylate kinase.* **Fig. 4d** illustrates AlphaFlex’s flexibility prediction (left panel) with the experimentally observed structural changes between its apo (green, PDB ID: 4akeB) and holo (blue, PDB ID: 2eckB) states (right panel). The major regions of structural rearrangement (highlighted by red arrows) show a strong correspondence with the regions AlphaFlex predicted as highly flexible, thereby demonstrating its effectiveness in identifying conformationally dynamic regions.

The performance of AlphaFlex to accurately identify flexible residues directly from an input structure suggests that static PDB databases, dynamic MD data, and structure- and sequence-derived features collaboratively encode rich, intrinsic information about protein conformational dynamics. Notably, our findings substantiate that the primary amino acid sequence fundamentally governs both protein tertiary structure and implicitly encodes crucial dynamic behavior information. This intrinsic information, in turn, shapes the energy landscape between multiple conformational states, enabling proteins to perform a series of functionally relevant conformational transitions under physiological conditions. This fundamental insight inspired our development of AlphaFlex, which implements selective masking of predicted flexible residue positions in MSA.

### Analysis of sampling performance

In this section, we perform ablation studies to analyze the effects of different model configurations, including MSA masking rate, depth-based sampling, and flexibility-guided masking, on the performance of generating multiple conformations. We speculate that the rich co-evolutionary coupling information within MSA not only underlies AlphaFold’s high-accuracy static structure predictions but also implicitly encodes dynamic information relevant to inter-conformational transitions^39^. Theoretically, these co-evolutionary patterns could reveal conformational heterogeneity and support modeling of multiple functional conformations. While AlphaFold achieves accuracy for single structure prediction, its default inference predominantly relies on a single, high-quality MSA, which inherently constrains its capacity to explore alternative or transitional states. To effectively decouple these dominant co-evolutionary constraints, we implemented column masking to the MSA based on predicted high flexibility residue regions, introducing gaps to weaken the original coupling information. This masking strategy was then combined with deep sampling to comprehensively explore the potential conformational space. Notably, the synergistic effect of MSA masking and deep sampling emerges as a critical factor enabling AlphaFold2 to generate multiple conformations.

To demonstrate the effectiveness of our MSA masking and sampling strategy in multiple conformation modeling, we designed three AlphaFlex variants for ablation analysis: w/o FG (without flexibility-guided MSA masking), w/o M (without MSA masking), and w/o S (without depth-based MSA subsampling), respectively. We evaluated these three variants together with the complete AlphaFlex method and the original AlphaFold2 (referred to as the Original baseline) on the benchmark dataset of 69 apo/holo protein pairs. We evaluated the average RMSD of the predicted structure relative to the apo and holo conformations across different ablation settings (**Figs 5a** and **5b**). Compared to the Original baseline, the w/o FG, w/o M, and w/o S variants each exhibited notable improvements. Specifically, w/o FG improved the average accuracy (RMSD) by 26.7% (apo) and 34.6% (holo), w/o M improved by 13.1% and 19.0%, and w/o S improved by 12.5% and 18.1%, respectively. These results highlight the importance of both MSA masking and subsampling in enhancing conformational accuracy.

**Fig. 5.**
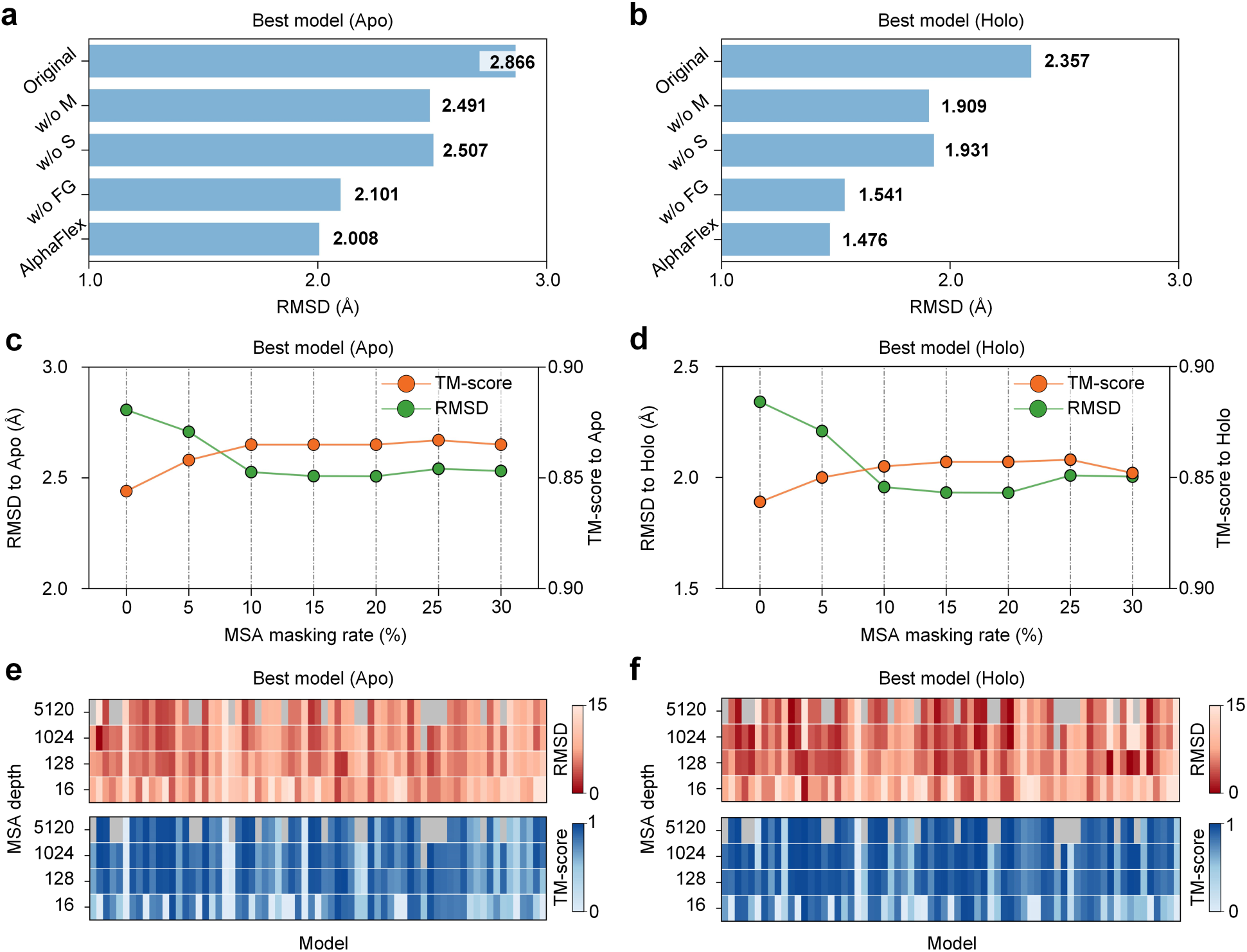
The impact of parameters on multiple conformation sampling performance. **a-b**, Average RMSD to the apo (**a**) and holo (**b**) states, comparing AlphaFold2 (Original), three ablated AlphaFlex variants (w/o FG: without flexibility guidance; w/o S: without deep subsampling; w/o M: without flexibility-guided masking), and the complete AlphaFlex method. **c-d,** Structural prediction accuracy (RMSD and TM-score) of AlphaFlex under varying guided masking ratios applied to flexible residues, evaluated separately for apo states (**c**) and holo states (**d**). **e-f,** Impact of MSA depth on prediction accuracy (RMSD and TM-score) across different protein targets for apo (**e**) and holo (**f**) states.

In order to further validate the effectiveness of our flexibility-guided MSA masking strategy, we designed the w/o FG variant by applying the same masking probability and subsampling strategy as AlphaFlex, but selecting the masking positions randomly (without using predicted flexible residue information). This experiment directly highlights the performance impacts of residue probability-guided MSA masking and random masking strategies. Compared to the AlphaFlex, the results show that without using the probability-guided masking strategy, prediction accuracy decreases by 4.4% for apo states and 4.2% for holo states. This demonstrates the effectiveness of our flexibility-guided MSA masking strategy in enabling AlphaFlex to capture high-accuracy conformational states.

MSA column masking guided by predicted flexible residue probability distribution represents a crucial determinant of AlphaFlex’s ability to generate accurate multiple conformations. Unlike random or large-scale masking methods, AlphaFlex applies this guided masking only to continuous fragments predicted to be highly flexible. This targeted strategy strategically relaxes evolutionary constraints while maintaining essential structural information, thereby facilitating the exploration of alternative conformational states. To assess the effect of different masking rates on structural prediction accuracy, we evaluated the performance using RMSD and TM-score metrics under guided masking ratios ranging from 0% to 30%. The detailed evaluation results are listed in **Supplementary Table 6**. For both apo and holo states (**Figs 5c** and **5d**), peak predictive performance was consistently achieved at a masking ratio of 20%, demonstrating an optimal balance between evolutionary constraint relaxation and structural information preservation. Instead of relying on large-scale random masking methods, which may lead to excessive information loss, we employed a sliding-window masking strategy that introduces perturbations in a controlled manner (see **Supplementary Fig. 1** for details). After evaluating the trade-off between prediction accuracy and computational cost, we implemented a 20% masking ratio for all subsequent analyses, while noting that protein-specific variations may require adjustment of this parameter.

MSA depth significantly influences the composition and expression of conformational information, which is crucial for exploring conformational space. To assess this influence on prediction accuracy, we analyzed prediction conformations under different MSA depths. Our analysis revealed that predicted model accuracy varies with MSA depth. We show the results for four representative MSA depths in **Figs. 5e** and **5f**, with full details for all depths provided in **Supplementary Fig. 2**. The accuracy of predicted structures for different protein targets generally improves with increasing MSA depth for both the apo and holo states. However, the optimal MSA depth required for high-accuracy prediction of conformations varies considerably across different proteins. Some proteins achieve high-quality predictions with relatively shallow MSAs, while others require much deeper alignments to achieve their best performance. This variability demonstrates the diverse evolutionary information inherent to different protein families and their unique conformational landscapes. These findings highlight the importance of strategically sampling MSAs at multiple depths to fully leverage co-evolutionary information for accurate and confident modeling of protein multiple conformations.

### Case studies

We have previously demonstrated that AlphaFlex can accurately model multiple conformations. In this section, we present case studies on two major categories: proteins that exhibit large-scale domain motions and those that show local loop motions. Through analyzing these cases, we have found that AlphaFlex not only recapitulates apo and holo states but also has the potential to capture intermediate transition states. Proteins undergo a series of conformational changes during the execution of their biological functions. However, the transient nature of these conformations and the complexity of the energy landscape pose significant challenges for structural modeling. Case studies on two typical types of protein, both exhibiting large-scale domain motions and local loop motions, show that AlphaFlex not only accurately reproduces their apo and holo states but also successfully predicts multiple continuous intermediate states between these two states.

Large-scale domain motions usually involve the relative rearrangement between two or more domains, accompanied by the overall rotation or translation of these domains (**Fig. 6a**), thereby causing significant conformational changes in proteins and driving the realization of their biological functions. For example, *adenylate kinase* (PDB ID: 4akeB) maintains intracellular metabolic balance by catalyzing the reversible reaction ATP+AMP↔2 ADP. Experimental evidence shows adenylate kinase undergoes substantial opening and closing of structural domains on a timescale of tens of microseconds, enabling its periodic catalytic cycle^40^. In the conformations generated by AlphaFlex, the RMSD relative to both apo and holo states shows a negative correlation (**Fig. 6a, i**), demonstrating that the methods not only accurately predict the two states but also generate continuous intermediate conformations from apo to holo and capture their conformational transition path. This smooth, nearly linear distribution indicates AlphaFlex’s high efficiency in exploring the intricate conformational space. It proficiently captures the subtle, dynamic changes characteristic of protein motions, offering crucial insights into transient intermediate states often elusive to experimental methods. In human *mitochondrial HSPD1* (PDB ID: 7azpA), the apical structural domain coordinates substrate binding and release via pronounced rotational motions and plays a crucial role in ensuring mitochondrial protein folding and homeostasis^41^. AlphaFlex accurately captures these rotational conformational transitions of the domain (**Fig. 6a, ii**). Similarly, in *DAHP synthase* (PDB ID: 1vr6A), binding of an aromatic allosteric inhibitor triggers a large-scale rotation of the apical domain, occluding the active site and achieving allosteric regulation^42^. AlphaFlex captures this inhibitor-induced conformational change (**Fig. 6a, iii**). Another example is the *D-ribose-binding protein* (PDB ID: 1urpD), which undergoes a lobe-closing motion upon ribose binding at the interdomain interface, securely clamping the ligand between the two lobes^43^. This conformational transition is also accurately captured by AlphaFlex (**Fig. 6a, iv**).

**Fig. 6.**
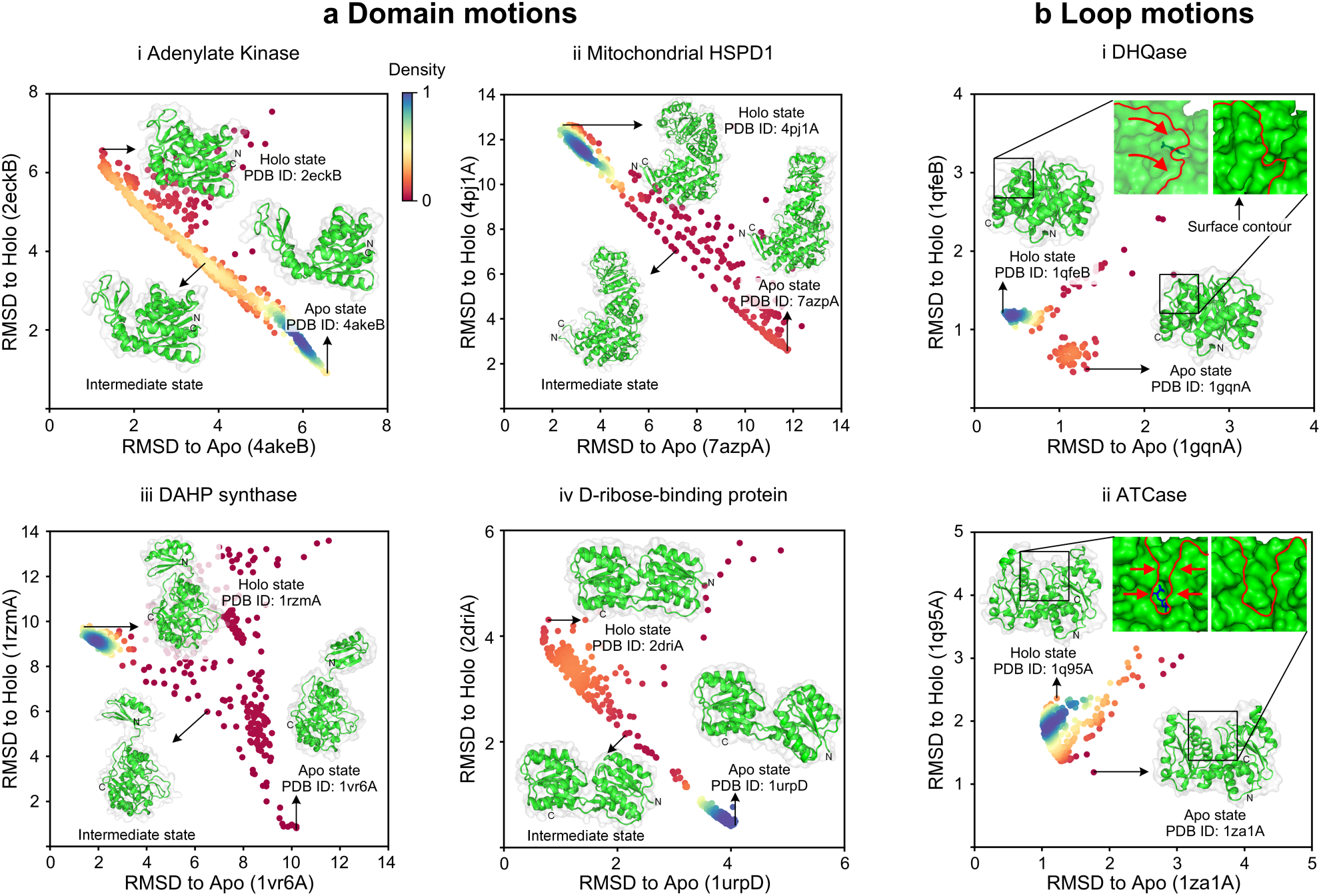
AlphaFlex predicts multiple conformational landscapes of functional proteins. **a**, Domain motions. Scatter plots characterize the conformational space sampled for proteins exhibiting significant domain movements, such as opening-closing or rotational motions. The x- and y-axes represent the RMSD from the experimentally resolved apo and holo states, respectively. Each point represents a predicted conformation, with its color indicating the conformational density (red represents the lowest density, while blue indicates the highest density). The color scale for density in each subplot is normalized from 0 to 1. Panels **i, ii, iii,** and **iv** present four representative examples of domain motion conformational changes generated by AlphaFlex. For these examples, the predicted best model is shown in green alongside the experimentally resolved apo and holo states outlined in white, as well as potential intermediate states. **b,** Loop motions. Similar scatter plots display conformational ensembles for proteins demonstrating dynamic loop movements, which are often involved in forming binding pockets. Insets highlight local surface contours where conformational changes occur.

Beyond large-scale domain motions, AlphaFlex also captures dynamic motions within loop regions (**Fig. 5b**). These motions are equally critical for enzymatic activity, ligand recognition, and gating mechanisms, often involving subtle conformational changes occurring on millisecond timescales that are challenging to resolve with conventional MD simulations. A representative case is *3-dehydroquinate dehydratase* (DHQase, PDB ID: 1gqnA), where a lid-like loop governs substrate access to the active site. Upon substrate binding, this lid-like loop closes to form a catalytically competent pocket, thereby facilitating the enzymatic reaction^44^ (**Fig. 5b, i**). Another illustrative example is *Aspartate Transcarbamoylase* (ATCase, PDB ID: 1za1A), where substrate binding follows a two-step induced fit mechanism. Initial binding leads to local electrostatic rearrangement within the active site, which subsequently triggers a cooperative domain closure^45^. This precise clamping motion stabilizes the substrate and promotes efficient catalysis. AlphaFlex not only recapitulates the open and closed states for these proteins but also accurately captures the conformational transitions between them, including the key loop motions involved (**Fig. 5b, ii**).

## Discussion

Although recent progress has been made in protein structure prediction, accurately capturing the multiple conformational states of proteins remains a significant challenge. Most existing models^1–4^ tend to predict a single static structure, thereby ignoring conformational heterogeneity and intermediate transition states. This limitation hinders a deeper understanding of the mechanisms underlying protein function. In this work, we propose AlphaFlex, a method for predicting multiple conformations. This method uses a deep learning network to extract inherent dynamic properties from protein homologous sequences and structures, enabling the prediction of residue flexibility probability distributions. These predicted distributions are then used to conditionally guide column masking in MSA, thereby achieving the modeling of multiple conformational states of proteins. The results demonstrate that AlphaFlex can effectively capture residual flexibility, leading to a notable improvement in multiple conformation prediction accuracy. The superior performance of AlphaFlex originates from its “flexibility-guided MSA processing” strategy.

By jointly predicting flexible residue probability distributions and applying deep subsampling to the same MSA, this approach effectively decouples co-evolutionary information contained within different conformations. As a result, AlphaFlex can explore a broader conformational space, thereby effectively addressing issues such as low sampling efficiency and poor conformational accuracy prevalent in existing methods. Despite the promising results, the applicability and accuracy of AlphaFlex could be further improved in several aspects. First, the MSAs employed in this study were generated from a limited set of sequence databases (see **Methods** for details) and may not capture rich homologous evolutionary information. Considering that the quality and diversity of MSAs are crucial factors influencing the performance of deep learning–based protein modeling, future efforts could benefit from incorporating more comprehensive sequence resources to construct deeper and more informative MSAs. Second, although the training set already includes structures derived from various experimental techniques, most of these structures predominantly represent stable conformations. This makes it challenging to fully reflect the conformational diversity and dynamic change proteins undergo under physiological conditions. To address this, future efforts could involve integrating data sources with richer conformational variation features, such as multi-state NMR structures, conformational ensembles resolved by cryo-EM, or high-quality MD simulation trajectories. This would enhance the model’s capacity to accurately model complex conformational transition pathways and transient intermediate states.

## Methods

Recent enhanced sampling methods^20–24^ based on AlphaFold2^1^ have demonstrated the ability to predict multiple protein conformations using subsampling strategies such as clustering, masking, or alanine substitution. Although these methods are effective and capable of exploiting co-evolutionary information, they usually rely on large random perturbations of the input information and do not effectively identify and exploit the intrinsic biological properties behind protein conformational changes, such as flexibility or dynamic regions. This lack of biological guidance in sampling presents a significant challenge in accurately and efficiently predicting multiple conformations.

To address this limitation, AlphaFlex introduces a biologically guided sampling framework that directs the decoupling of co-evolutionary constraints specifically within flexible regions, rather than relying on random perturbations. The prediction of multiple conformations is formulated as a two-step process. First, AlphaFlex accurately predicts residue flexibility probability distribution. This prediction provides a crucial probabilistic map of potential dynamic regions within the protein. Second, we use the predicted flexibility probabilities to perform directional masking on the MSA, coupled with deep sampling to generate distinct sub-MSAs. This strategic masking effectively decouples co-evolutionary constraints within flexible regions, thereby prompting AlphaFold2 to explore and generate diverse and plausible conformations.

### Construction of multiple conformation datasets

AlphaFlex was initially pre-trained on the static structures from the Protein Data Bank (PDB)^33^, and performed fine-tuned on dynamic conformations from the MD simulation database ATLAS^46^. The workflow of the dataset construction process is provided in **Supplementary Fig. 3**. The steps for dataset construction are as follows:

1. We first collected all proteins released before January 1, 2024, including monomeric and complex structures determined by various experimental techniques such as X-ray diffraction, nuclear magnetic resonance (NMR), and cryo-electron microscopy (cryo-EM). After splitting all complexes into monomers, a total of 608,984 monomer structures were obtained.
2. All protein sequences were clustered using MMseqs^47^ at an 80% similarity threshold, resulting in 65,327 clusters. The cluster centers were selected as representative structures and sequences.
3. Clusters with representative sequences between 50 and 500 residues were retained and subsequently filtered for redundancy at a 30% sequence similarity threshold, yielding 20,932 non-redundant clusters. Ultimately, we utilized a training set comprising 17,485 clusters released before January 1, 2023, and the remaining 1,937 classes as a validation set.
4. To fine-tune our model, we selected 1,390 non-membrane proteins from the ATLAS^46^ dataset. For each protein, 100 frames were uniformly sampled from the first of the three available MD trajectories to construct conformational clusters. After removing clusters with sequences shorter than 50 or longer than 500 amino acids, 1,279 clusters were retained. These clusters were then partitioned into training (1,172) and validation (107) sets based on their release dates, with 1 May 2018 as the cutoff date.

### Test set

To evaluate the model’s performance in predicting flexible residues and multiple conformation predictions, we used a widely recognized benchmark dataset comprising 91 apo/holo proteins, originally derived from the CoDNaS^27^ database. This dataset is commonly employed for assessing protein conformational change prediction. Each protein in this dataset consisted of two experimentally determined structures, representing its apo and holo conformational states. All structure pairs underwent manual curation to ensure that the observed conformational changes were supported by experimental evidence and associated with biological functions^49^, rather than caused by artifacts, alignment errors, unresolved regions, or flexible termini. Furthermore, for proteins with more than two conformers, the pair exhibiting the largest RMSD between the apo and holo structures was selected to represent the principal conformational change. Other selection criteria included the absence of intrinsic disorder, high-resolution crystal structures, and no sequence discrepancies or mutations. To ensure the independence of the evaluation set from the training set, we performed redundancy reduction using MMseqs2 with a 30% sequence identity cutoff, resulting in a final test set of 69 apo/holo proteins, with an average RMSD of 3.6Å between their apo and holo states. The names of all test proteins are listed in **Supplementary Table 7**, with the RMSD and TM-score between apo and holo protein pairs provided in **Supplementary Table 8**. The distribution of RMSD and TM-score values between apo and holo protein pairs is provided in **Supplementary Fig. 4**.

We further evaluated AlphaFlex’s performance in multiple conformation predictions on a membrane protein test set^38^. This test set initially comprised 16 membrane proteins, each with experimentally determined inward-facing and outward-facing conformations. To ensure the independence of this evaluation set from the training set, we performed redundancy reduction using MMseqs2 with a 30% sequence identity cutoff, resulting in a final test set of 5 proteins. The names of all test membrane proteins are provided in **Supplementary Table 9**, with the RMSD and TM-score between apo and holo protein pairs provided in **Supplementary Table 10.**

### Input features

Our deep learning model for per-residue flexibility prediction integrates diverse input features derived from an initial protein structural input. These features are designed to provide a comprehensive representation of each residue across its sequence, structural environment, MSA context, and structure profile information. Specifically, these features include one-hot encoding, the BLOSUM62 substitution matrix, Ultrafast Shape Recognition (USR), distance and orientation matrices, physicochemical descriptors, MSA embedding vectors obtained from a protein language model (ESM-MSA-1b)^50^, and structural profile features obtained from homologous structures. Together, they provide a rich representation of each residue across sequence, structure, and MSA. Detailed descriptions of all input features are provided in **Supplementary Table 1**.

### Model architecture

We developed a structure-informed deep learning model to predict per-residue flexibility probability distributions. The model’s network architecture comprises a feature integration module, an attention mechanism, and a deep residual network. Initially, structural features (geometric and physicochemical properties) and structural profile are fused within the feature integration module to capture 3D structural contextual information. Simultaneously, MSA embeddings are linearly projected and layer-normalized before being processed through a sequence reweighting module that assigns weights to each MSA sequence, extracting co-evolutionary features per residue. All processed features are concatenated and fed into an attention module composed of three layers of triangle updates and axial attention, capturing long-range dependencies and contextual interactions. The output of the attention module is fed into a 64-layer deep residual network. Each residual block consists of convolutional layers, instance normalization, and activation functions. Specifically, each residual block contains two 3×3 convolutional layers, each followed by instance normalization and ELU activation. Finally, the output is processed through a fully connected layer followed by a sigmoid activation to produce the predicted flexibility score for each residue.

### Training

Collecting sufficient MD simulation data to comprehensively characterize protein conformational changes remains a major challenge, primarily due to the limited data availability and the high degree of diversity and heterogeneity^46^. To address this data scarcity, we adopted a transfer learning strategy. AlphaFlex was first pre-trained on our self-constructed protein multiple conformation dataset. Subsequently, based on the best-performing model from the pre-training model, we fine-tuned it using the ATLAS MD dataset to further capture protein dynamic information.

All neural networks were implemented in Python using the PyTorch framework. Training was performed using the Adam optimizer with an initial learning rate of 1e-4 and a decay rate of 0.99. The Focal Loss function was employed for optimization and is define as follows:

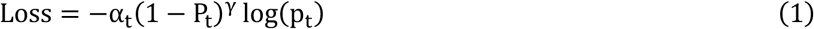

where *p*_t_ denotes the predicted probability for the positive class, *a*_t_ is the balancing factor for class imbalance, and γ is the focusing parameter that down-weights well-classified samples. In this study, α was set to 0.25 and γ to 2.

The model with the lowest validation loss during training was selected for downstream tasks. All training was carried out on a machine equipped with one NVIDIA A100 GPU and two Intel Xeon(R) Gold 6248 processors, with the total training cost summarized in **Table 1**.

**Table 1.** AlphaFlex training cost.

| Training phase | Number of samples | Total hours | Hours per epoch |
| --- | --- | --- | --- |
| Pre-training (PDB) | 165k | 64 | 2.5 |
| Fine-tuning (MD) | 118k | 11 | 0.5 |

### MSA generation

To obtain more diverse MSAs, we used the JackHMMR algorithm^51^ to query sequences from the UniRef90^52^, and the HHblits algorithm^55^ to query sequences from the UniRef30^56^ database and BFD^53,54^ databases, to obtain the MSAs for each protein. Detailed descriptions of the above databases are provided in **Supplementary Note 2**.

### MSA masking strategy

To effectively decouple protein dynamic information while preserving crucial co-evolutionary information, we applied a sliding window progressive masking strategy on predicted high-probability flexible residue regions. For each identified flexible region, a sliding window of size 3 was moved stepwise by 1 residue from both ends toward the center for masking. Detailed descriptions of the masking strategy are provided in **Supplementary Fig. 1**. The resulting masked sub-MSAs were then deep-sampled at sequence depths of 16, 32, 64, 128, 256, 512, 1024, 1536, 2048, 2560, 3072, 3584, 4096, 4608, and 5120. The final sub-MSAs generated in this way were used for downstream analyses.

### Conformation generation

We used the processed sub-MSAs as input to generate protein structures with AlphaFold^1^ (available at https://github.com/google-deepmind/alphafold), without using templates in this process.

### Baseline approaches

A range of baseline approaches was selected for experimental analysis. AF-Cluster was run with the default parameters as described in its GitHub repository (https://github.com/HWaymentSteele/AF_Cluster). For AF2_conformations, we randomly set max_extra_msa to any of these values: [16, 32, 64, 128, 256, 512, 1024, 5120], and max_msa_clusters was set either half of max_extra_msa or a maximum of 512, and the code is provided at https://github.com/delalamo/af2_conformations. AlphaFLOW was run with its default parameters as described in its GitHub repository (https://github.com/bjing2016/alphaflow). Across all methods, a total of 1000 conformations were generated per target.

### Performance evaluation metrics

In this study, we employed multiple metrics to evaluate the performance of the flexible residue prediction model, including AUC-ROC, accuracy, precision, and F1-score. Additionally, we used success rate, RMSD, and TM-score to quantify the proximity between AlphaFlex-generated models and experimental structures. The success rate was defined as the proportion of targets for which both the predicted apo and holo structures achieved TM-score greater than 0.95, indicating high structural similarity to the respective experimental states. Both RMSD and TM-score metrics were calculated using the TM-score^36^ tool available at https://zhanggroup.org/TM-score/. Detailed descriptions of the above metrics are provided in **Supplementary Note 1**.

## Data availability

The data that support this study are available from the corresponding authors upon request.

## Code availability

Release of inference code and model weights is in preparation.

## Acknowledgements

We thank members of the Guijun Zhang lab for discussion and feedback; X.Chen for helping with dataset preparations. Computational resources were provided by the College of Information Engineering at Zhejiang University of Technology. This work was supported by the National Key R&D Program of China [2022ZD0115103], the National Nature Science Foundation of China (62173304, 62203389), the “Pioneer” and “Leading Goose” R&D Program of Zhejiang (2025C01190), and the Zhejiang Province High-level Talent Special Support Program (2023R5248).

## Author contributions

G.Z. conceived and supervised the research. Y.Z. helped supervise the research. G.Z., L.G., and X.Cui designed the experiment. G.Z., L.G., and X.Cui collected the data and performed the experiment. G.Z., L.G., X.Cui, K.Z., Y.Z., and X.Z. analyzed the data. G.Z., L.G., and X.Cui wrote the manuscript, and all authors read and approved the final manuscript.

## Additional information

Correspondence and requests for materials should be addressed to Guijun Zhang.

## Supplementary Information for

### Supplementary Notes

#### Supplementary Note 1. Description of evaluation metrics

##### RMSD

Root-mean-square deviation (RMSD) is a metric for quantifying the average distance between a set of corresponding atoms in two protein structures. It is calculated by

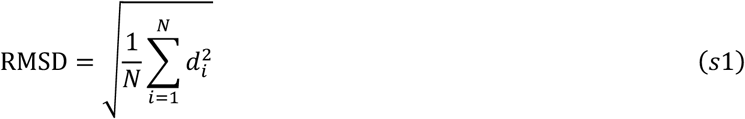

where *N* is the number of corresponding atoms (C_α_) in the two structures, and *d*_i_ is the Euclidean distance between the *i*-th pair of corresponding atoms after optimal superposition (translation and rotation) of one structure onto the other. The goal of superposition is to minimize the RMSD value. The value of RMSD is non-negative and typically expressed in Angstroms (Å). A lower RMSD value indicates greater structural similarity between the two proteins. While widely used, RMSD can be sensitive to local structural differences and outliers, particularly in flexible or disordered regions.

##### TM-score

TM-score^1^ is a metric for evaluating the topological similarity between protein structures, which can be calculated by

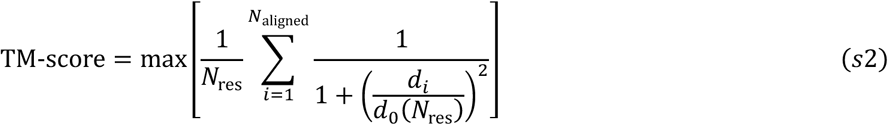

where *N*_res_ is the amino acid sequence length of the target protein, *N*_aligned_ is the length of the aligned residues to the reference (native) structure, *d*_i_ is the distance between the *i*-th pair of aligned residues, *d_0_(N_res_ = 1.24^3^√N_res_ − 15 − 1.8* is a scale to normalize the match difference, and ‘max’ refers to the optimized value selected from various rotation and translation matrices for structure superposition. The value of TM-score ranges in (0,1], where a higher value indicates closer structural similarity. Stringent statistics showed that TM-score >0.5 corresponds to a similarity with two structures having the same fold and/or domain orientations^2^.

##### Success Rate

The success rate is defined as the proportion of targets for which both the predicted apo and holo conformations achieve TM-scores greater than 0.95. It is calculated as the number of such successful cases divided by the total number of targets. It is calculated by

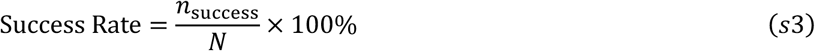

where *N* is the number of protein targets, *n*_success_ is the number of targets where both the predicted apo and holo conformations have TM-score^1^ > 0.95.

#### Supplementary Note 2. Description of the databases

##### UniRef30

The UniRef30^3^ database is an HHblits-style^4^ hidden Markov model (HMM) database that clusters UniProtKB^5^ sequences at the level of 30% pairwise sequence identity by MMseqs2^6^. For each cluster, an “A3M” formatted MSA generated by Clustal Omega^7^ and the corresponding HMM are provided for HHblits. UniRef30 is the vision of the database generated after 2019. In total, UniRef30 provides 231 million sequences in 25 million clusters.

##### Uniref90

Uniref90^8^ provides sequences from the UniProtKB^8^ clustered at 90% pairwise sequence identity by MMseqs2. Unlike Uniclust30 and UniRef30 (which are HMM databases), Uniref90 is a flat sequence database. For each cluster, only the representative sequence of the cluster is kept in the database. Thus, there are 109 million protein sequences/ clusters in the Uniref90 database.

##### BFD

The BFD is an HHblits-style HMM database that was created by clustering 2.5 billion protein sequences from UniProtKB^8^, Metaclust^9^, the soil reference catalog, and the marine eukaryotic reference catalog assembled by Plass^10^. BFD was clustered by MMseqs2^6^ with 30% pairwise sequence identity, and only the clusters that had more than three sequence members were kept in the database. In total, 66 million clusters and 2.2 billion genomics/metagenomics sequences are collected in the BFD database.

## Supplementary Tables

**Supplementary Table 1.** Detailed description of input features for the network.

| Feature |  | Description | Shape/Dimensions |
| --- | --- | --- | --- |
| Geometry | One-hot encoding | Represents each amino acid in the sequence as binary vectors, where a single “1” indicates the identity of the amino acid at each position. | L×20 |
|  | Ultra-fast Shape Recognition (USR) | Able to quickly capture the relationship between local and global topological structures | L×3 |
|  | Secondary structures | One-hot encoded representation of three state secondary structures given by DSSP solver. | L×4 |
|  | Voxelization | Voxelization is a method of converting a protein's 3D structure into a 3D grid (voxels) representation. The atomic distribution or properties of the protein are mapped onto discrete voxel units, forming a 3D tensor. | L×640 |
|  | Distance map | Represents the spatial distances between all pairs of amino acid residues in a protein. | L×L×4 |
|  | Orientation map | Represents the spatial orientation information between pairs of residues in a protein. | L×L×6 |
| Evolutionary | BLOSUM62 Substitution Matrix | Encodes evolutionary and functional similarity between residues using a substitution matrix. Represents L×24 replacement vectors for each residue based on its position in the sequence. | L×24 |
| Physicochemical | One-body energy term | p_aa_pp, rama_prepro, omega, fa_dun. | L×L×7 |
|  | Two-body energy term | fa_atr, fa_rep, fa_sol, lk_ball_wtd, fa_elec, hbond_bb_sc, and hbond_sc. | L×7 |
| MSA Embeddings | Attention Matrix | Encapsulates deep contextualized features of each residue generated by the pre-trained ESM-MSA-1b language model. Includes evolutionary, structural, and functional information. | L×L×144 |
|  | Feature Matrix |  | L×768 |
| Structure profile | Structure profile | Characterizing the potential spatial displacement distance of each residue from a profile constructed from structure clusters. | L×1 |

**Supplementary Table 2.**
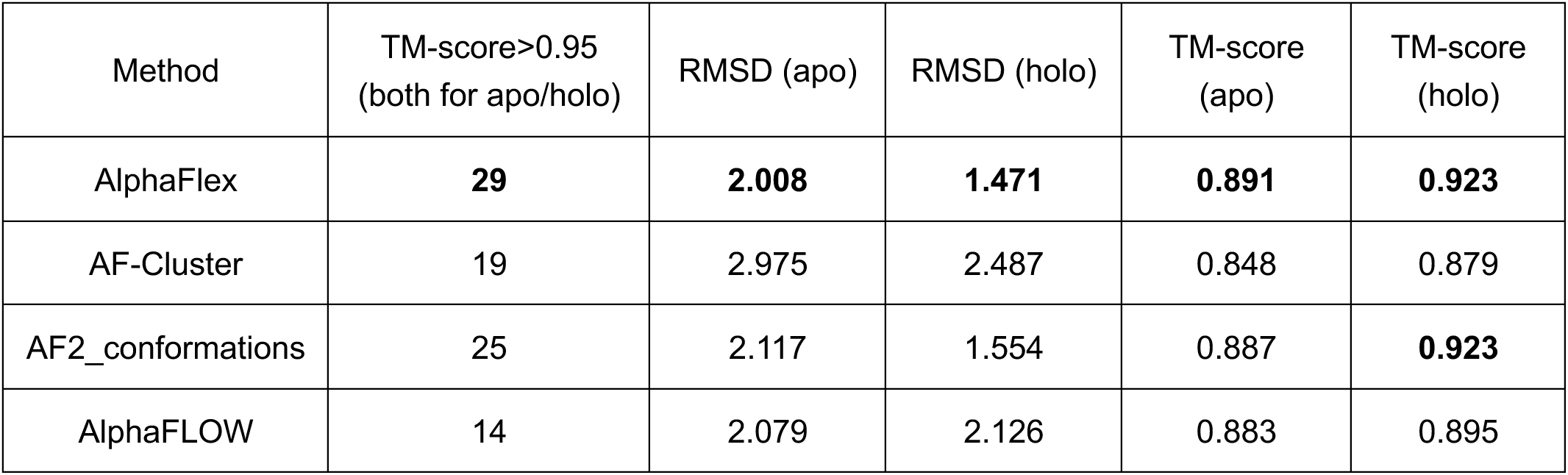
Analysis of the best multiple conformations of AlphaFlex, AF-Cluster, AF2_conformations, and AlphaFLOW on 69 apo-holo protein pairs.

| Method | TM-score>0.95<br>(both for apo/holo) | RMSD (apo) | RMSD (holo) | TM-score<br>(apo) | TM-score<br>(holo) |
| --- | --- | --- | --- | --- | --- |
| AlphaFlex | **29** | **2.008** | **1.471** | **0.891** | **0.923** |
| AF-Cluster | 19 | 2.975 | 2.487 | 0.848 | 0.879 |
| AF2_conformations | 25 | 2.117 | 1.554 | 0.887 | **0.923** |
| AlphaFLOW | 14 | 2.079 | 2.126 | 0.883 | 0.895 |

**Supplementary Table 3.** Analysis of the best multiple conformations of AlphaFlex, AF-Cluster, AF2_conformations, and AlphaFLOW on 5 membrane proteins.

| Method | RMSD<br>(inward-facing) | RMSD<br>(outward-facing) | TM-score<br>(inward-facing) | TM-score<br>(outward-facing) |
| --- | --- | --- | --- | --- |
| AlphaFlex | **2.716** | **3.083** | **0.912** | **0.887** |
| AF-Cluster | 4.134 | 3.528 | 0.851 | 0.865 |
| AF2_conformations | 2.941 | 3.389 | 0.900 | 0.868 |
| AlphaFLOW | 4.038 | 3.104 | 0.847 | 0.884 |

**Supplementary Table 4.** Comparison of flexible residue prediction performance by different methods using ROC-AUC, Accuracy, Precision, and F1 score metrics.

| Method | ROC-AUC | Accuracy | Precision | F1 score |
| --- | --- | --- | --- | --- |
| AlphaFlex | **0.810** | **0.742** | **0.491** | **0.521** |
| AF-Cluster | 0.557 | 0.423 | 0.300 | 0.402 |
| AF2_conformations | 0.564 | 0.491 | 0.342 | 0.368 |
| AlphaFLOW | 0.658 | 0.720 | 0.442 | 0.424 |

**Supplementary Table 5.** Ablation study on the impact of different features in AlphaFlex for flexible residue prediction.

| Method | ROC-AUC | Accuracy | Precision | F1 score |
| --- | --- | --- | --- | --- |
| AlphaFlex | **0.810** | **0.742** | **0.491** | **0.521** |
| w/o Geometry | 0.792 | 0.712 | 0.452 | 0.520 |
| w/o Evolutionary | 0.807 | 0.718 | 0.470 | 0.499 |
| w/o Physicochemical | 0.784 | 0.738 | 0.473 | 0.489 |
| w/o MSA embeddings | 0.779 | 0.696 | 0.446 | 0.491 |
| w/o Structure profile | 0.806 | 0.718 | 0.461 | 0.492 |

**Supplementary Table 6.** Evaluation of AlphaFlex performance (RMSD, TM-score) across different masking rates of multiple conformations in apo and holo states.

| MSA masking (%) | RMSD (apo) | RMSD (holo) | TM-score (apo) | TM-score (holo) |
| --- | --- | --- | --- | --- |
| 0 | 2.807 | 2.341 | 0.844 | 0.889 |
| 5 | 2.708 | 2.209 | 0.858 | 0.900 |
| 10 | 2.526 | 1.957 | 0.865 | 0.905 |
| 15 | 2.508 | 1.932 | 0.865 | 0.907 |
| 20 | **2.507** | **1.931** | 0.865 | 0.907 |
| 25 | 2.541 | 2.009 | **0.867** | **0.908** |
| 30 | 5.531 | 2.004 | 0.865 | 0.902 |

**Supplementary Table 7.** The 69 targets of the protein test set.

| No. | Apo | Holo | No. | Apo | Holo | No. | Apo | Holo |
| --- | --- | --- | --- | --- | --- | --- | --- | --- |
| 1 | 1AEL-12_A | 1URE-8_A | 24 | 1K6W_A | 1K70_A | 47 | 1W4U-9_A | 1UR6-4_A |
| 2 | 1AKZ_A | 1SSP_E | 25 | 1L0W_B | 1G51_A | 48 | 1XSA-23_A | 1XSC-20_A |
| 3 | 1AW2_A | 1AW1_B | 26 | 1LIP-2_A | 1JTB-6_A | 49 | 1ZA1_A | 1Q95_A |
| 4 | 1BV2-3_A | 1UVB_A | 27 | 1LMZ-9_A | 1P7M-18_A | 50 | 2AI6-18_A | 2OZW-3_A |
| 5 | 1C54-19_A | 1RGH_B | 28 | 1MO7-3_A | 1MO8-14_A | 51 | 2BNH_A | 1DFJ_I |
| 6 | 1DMO-18_A | 3CLN_A | 29 | 1MUT-11_A | 1PUN-7_A | 52 | 2CJO-5_A | 1ROE-10_A |
| 7 | 1E5L_A | 1E5Q_A | 30 | 1MX7-20_A | 1MX8-25_A | 53 | 2D9E-12_A | 2RS9-13_B |
| 8 | 1EAL-2_A | 1EIO-3_A | 31 | 1NJG_B | 1NJF_A | 54 | 2F63-4_A | 1EQM_A |
| 9 | 1EVK_A | 1EVL_A | 32 | 1O1U-9_A | 1O1V-5_A | 55 | 2FHM-12_A | 2HLT-19_A |
| 10 | 1F3Y-17_A | 1JKN-15_A | 33 | 1ORM-9_A | 1QJ8_A | 56 | 2IN2_A | 2B0F_A |
| 11 | 1FSF_A | 1FQO_B | 34 | 1OTJ_D | 1GY9_A | 57 | 2JU3-3_A | 2JU8-9_A |
| 12 | 1G6W_D | 1K0B_C | 35 | 1PDB_A | 1YHO-9_A | 58 | 2JWW-13_A | 1RTP_1 |
| 13 | 1GQN_A | 1QFE_B | 36 | 1PFL-19_A | 1FIL_A | 59 | 2K43-1_A | 2K8R-1_A |
| 14 | 1GUD_A | 1RPJ_A | 37 | 1RF5_A | 1RF4_A | 60 | 2KXL-8_A | 2K0G-9_A |
| 15 | 1HOO_B | 1CG0_A | 38 | 1RKA_A | 1GQT_A | 61 | 2L68-9_A | 2LKK-10_A |
| 16 | 1HW1_B | 1H9G_A | 39 | 1RKM_A | 1B6H_A | 62 | 2LAO_A | 1LAH_E |
| 17 | 1I56_A | 1EL1_A | 40 | 1S2O_A | 1TJ5_A | 63 | 2LHS-6_A | 2BEM_A |
| 18 | 1IGP_A | 2AU6_A | 41 | 1SKT-10_A | 1TNQ-33_A | 64 | 2LKC-4_A | 2LKD-18_A |
| 19 | 1IJA-15_A | 2KID-4_A | 42 | 1SYM-2_B | 1XYD-15_B | 65 | 2NLN-4_A | 1RRO_A |
| 20 | 1JEJ_A | 1JG6_A | 43 | 1TFU_A | 3UC5_A | 66 | 2UZ5-9_A | 2VCD-8_A |
| 21 | 1JFJ-3_A | 1JFK_A | 44 | 1URP_D | 2DRI_A | 67 | 4AKE_B | 2ECK_B |
| 22 | 1K2H-4_A | 1ZFS-13_B | 45 | 1VIY_C | 1VHL_A | 68 | 6OY9_B | 1A0R_B |
| 23 | 1K5H_A | 1Q0Q_A | 46 | 1VR6_A | 1RZM_A | 69 | 7AZP_A | 4PJ1_A |

**Supplementary Table 8.**
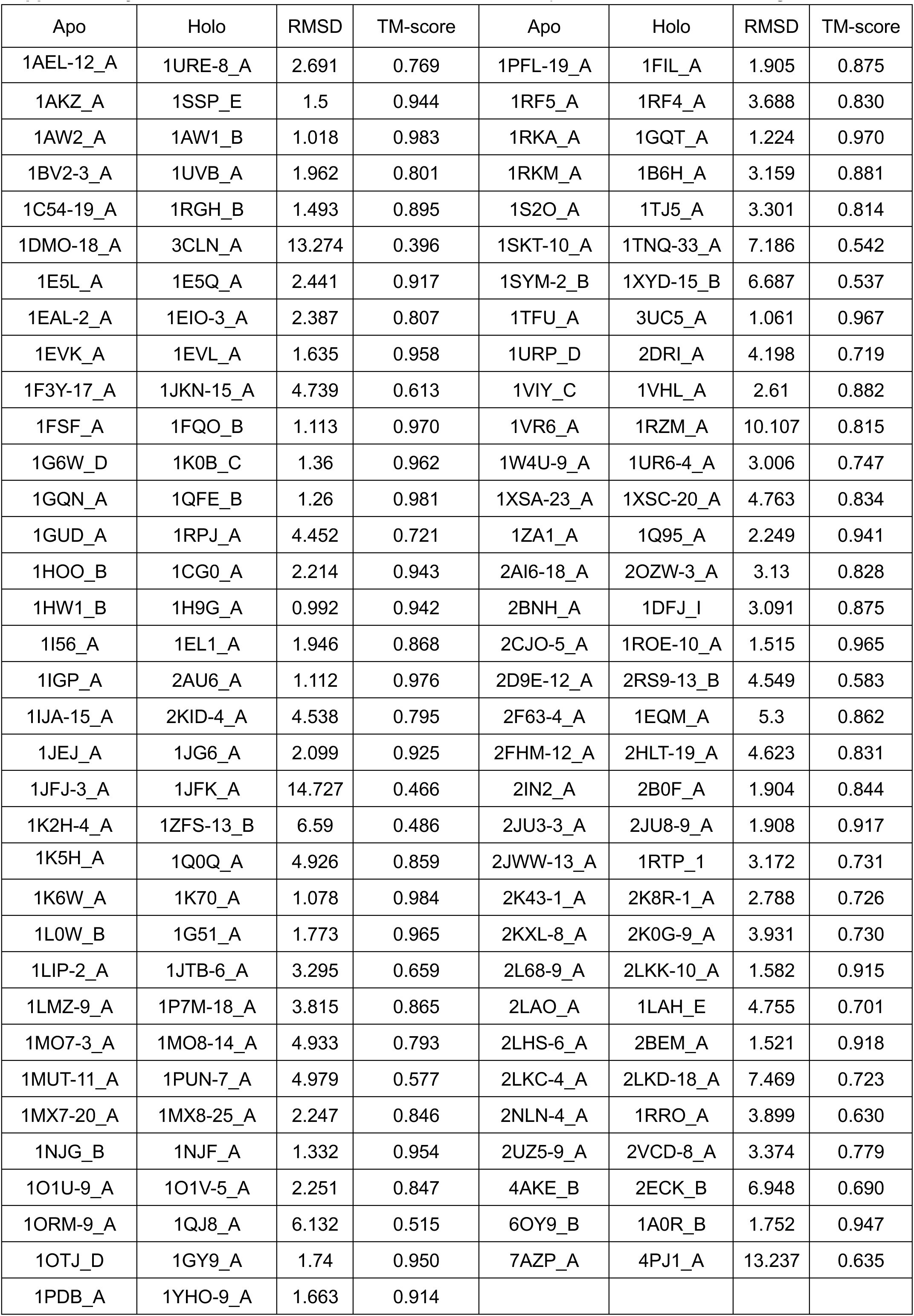
The RMSD and TM-score between the apo and holo states of 69 targets.

**Supplementary Table 9.** The 5 targets of the Membrane protein test set.

| No. | Inward-facing conformation | Outward-facing conformation |
| --- | --- | --- |
| 1 | 3QF4_A | 6QV1_C |
| 2 | 7L17_A | 4M61_A |
| 3 | 6BVG_A | 5IWS_A |
| 4 | 6VS1_A | 6GV1_A |
| 5 | 6NC7_A | 6NC9_A |

**Supplementary Table 10.** The RMSD and TM-score between the inward-facing and outward-facing conformations of 5 targets.

| Inward-facing conformation | Outward-facing conformation | RMSD | TM-score |
| --- | --- | --- | --- |
| 3QF4_A | 6QV1_C | 5.606 | 0.740 |
| 7L17_A | 4M61_A | 5.738 | 0.746 |
| 6BVG_A | 5IWS_A | 7.313 | 0.671 |
| 6VS1_A | 6GV1_A | 4.583 | 0.758 |
| 6NC7_A | 6NC9_A | 6.817 | 0.650 |

## Supplementary Figures

**Supplementary Fig. 1.**
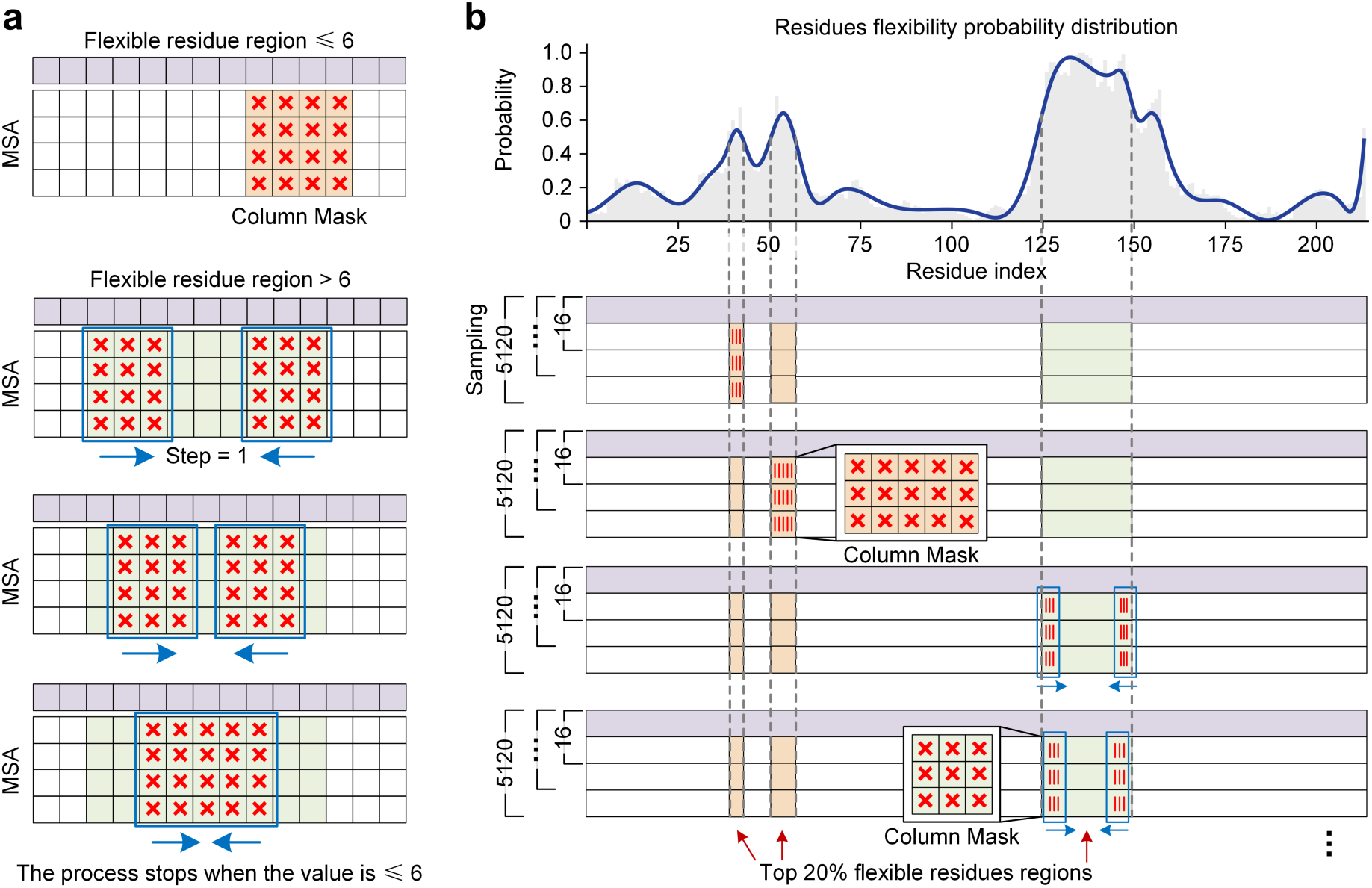
MSA Masking Strategy. a, If the flexible residue region is ≤6 amino acids, apply direct column masking. If the flexible residue region is >6 amino acids, two sliding windows of size 3 are simultaneously applied from both sides of the region, moving towards each other with a step size of 1 until they meet. b, We first identified the top 20% residues with the highest predicted flexibility probability from the distribution map, then applied different masking strategies to these regions according to the length of the amino acid region, applied a deep sampling strategy, and finally generated sub-MSA.

**Supplementary Fig. 2.**
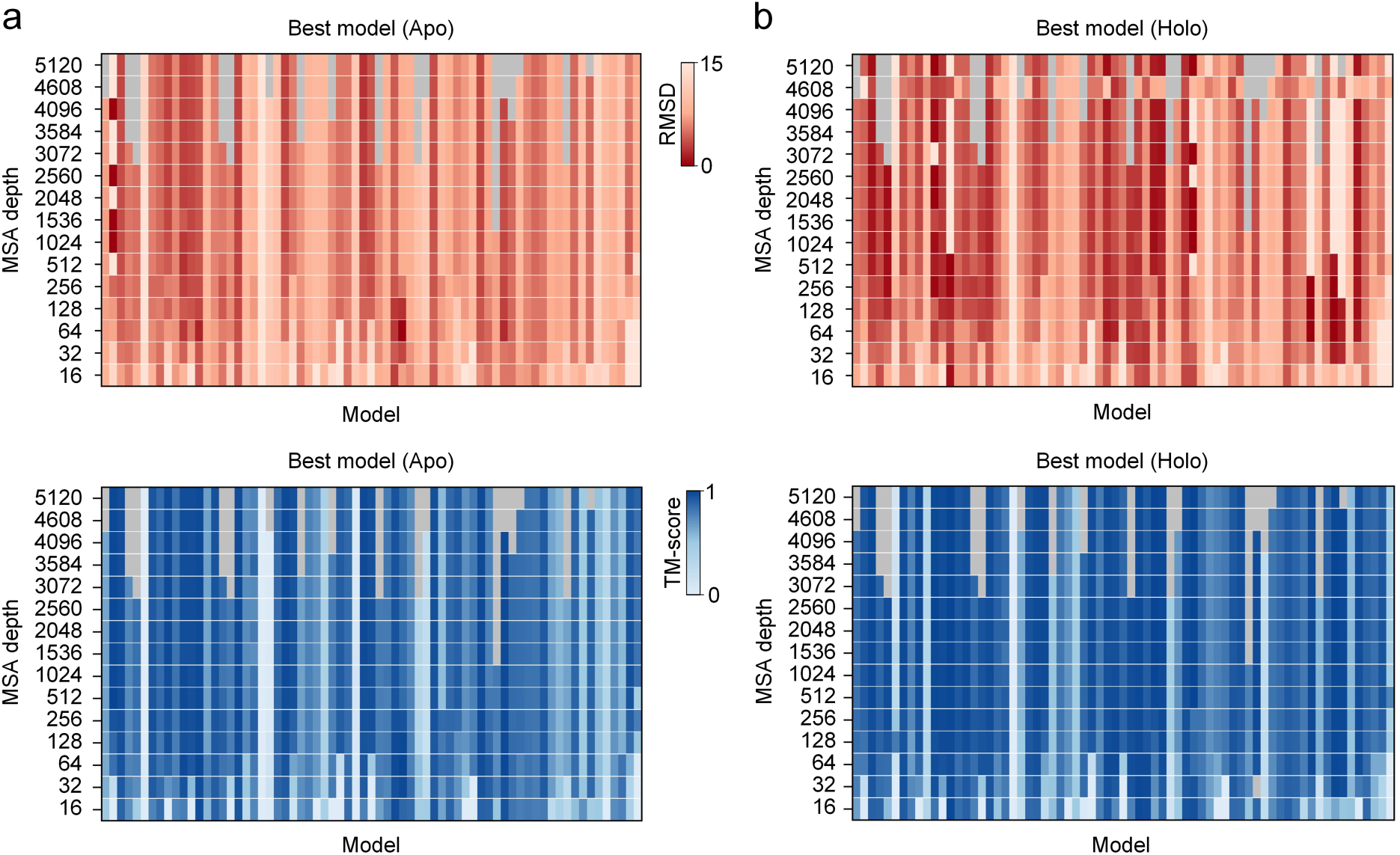
The impact of MSA depth parameters on multiple conformation sampling performance. a-b, Impact of MSA depth on prediction accuracy (RMSD and TM-score) across different protein targets for apo (a) and holo (b) states.

**Supplementary Fig. 3.**
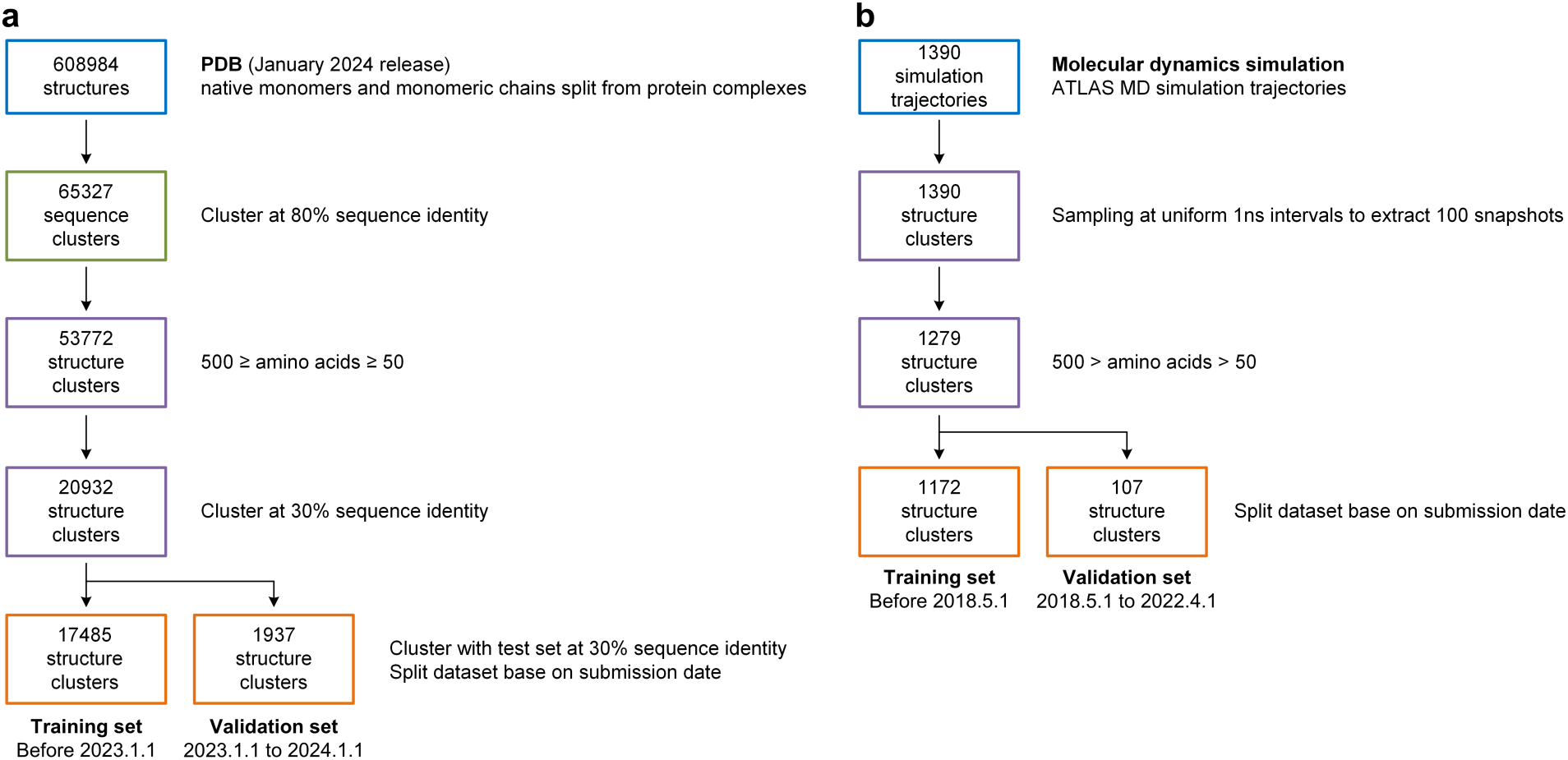
Construction of multiple conformation training datasets. **a**, PDB^11^ structures (January 2024 release), including native monomers and monomeric chains split from protein complexes, were initially collected. These structures were then clustered at 80% sequence identity, resulting in 65,327 sequence clusters. From these clusters, structures with an amino acid length between 50 and 500 residues were selected, yielding 53,772 structure clusters. These clusters were further clustered at 30% sequence identity, reducing the count to 20,932 structure clusters. Further clustering with the test set at 30% sequence identity was performed, and the dataset was divided into training and validation sets based on the submission date: 17,485 structures submitted before January 1, 2023 were designated as the training set, and 1,937 structures submitted between January 1, 2023 and January 1, 2024 constituted the validation set. **b**, For molecular dynamics (MD) data, 1390 simulation trajectories were collected from the ATLAS^12^ database. These trajectories were then processed by sampling at uniform 1ns intervals to extract 100 snapshots, resulting in 1390 initial structure clusters. Subsequently, structure clusters with amino acid lengths between 50 and 500 residues were retained, reducing the set to 1279 clusters. The dataset was then partitioned based on submission date: 1172 structure clusters submitted before May 1, 2018, were designated as the training set, and 107 structure clusters submitted between May 1, 2018, and April 1, 2022, formed the validation set.

**Supplementary Fig. 4.**
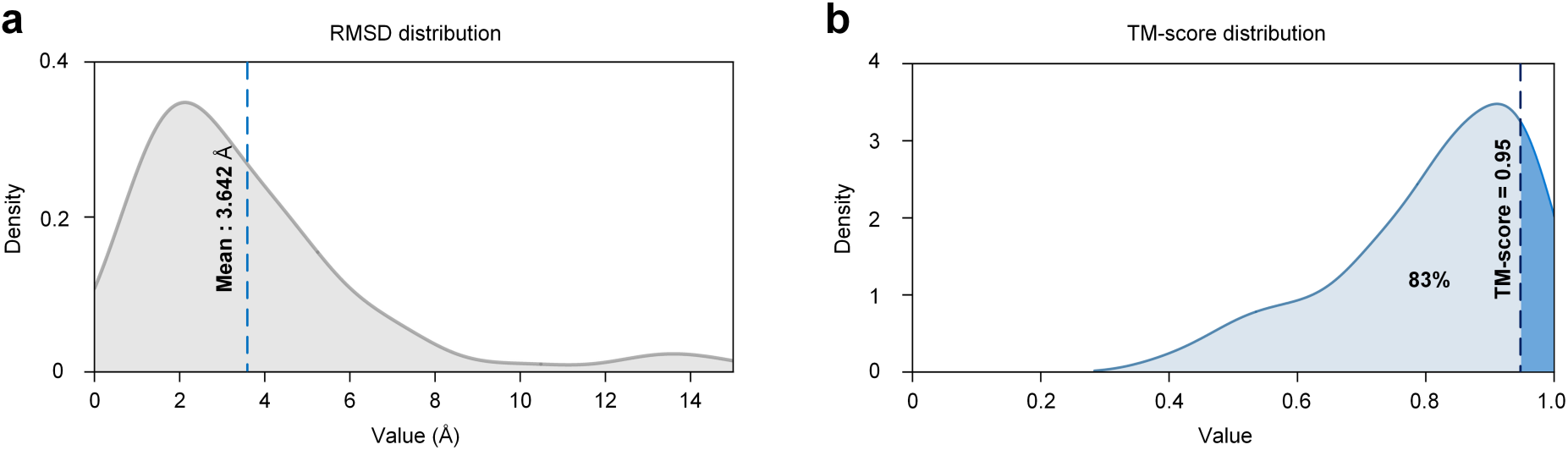
Construction of multiple conformation training datasets. a, The distribution of RMSD and TM-score values between apo and holo protein pairs. b, The average RMSD between the two states is 3.364Å. 83% of the target two states have a TM-score less than 0.95.

